## Supplemental Material for "Axonemal regulation by curvature explains sperm flagellar waveform modulation"

July 20, 2022

### 1 Numerical discretisation

As highlighted in the main manuscript, the full system of equations for the ARCH model: Eqs. (1-2), and Eqs. (5-8) are discretised following the approach contained within Refs. [1, 2], with the non-local hydrodynamics accounted for using the parallelised NEAREST method of Smith and Gallagher [3, 4, 5, 6].

#### 1.1 Head discretisation

The sperm head is modelled as an ellipsoidal surface, semi-axes lengths ($2\ \mu\text{m}, 1.6\ \mu\text{m}, 1\ \mu\text{m}$), inclined at an angle $\phi(t)$ to the horizontal, and discretised with $n_H = 24$ vector degrees of freedom (DOF) for the coarse force discretisation, denoted $\mathbf{Y}^{[n]}$ for $n = 1, \ldots, n_H$, and $n_Q = 384$ DOF for the fine quadrature discretisation, denoted $\hat{\mathbf{Y}}^{[n]}$ for $n = 1, \ldots, n_Q$. The force per unit area exerted by the head onto the fluid is denoted $\boldsymbol{\varphi}^{[n]} := \boldsymbol{\varphi}(\mathbf{Y}^{[n]}, t)$, $n = 1, \ldots, n_H$.

#### 1.2 Flagellum discretisation

The flagellum is discretised into $n_T = 60$ straight-line segments of equal length $\Delta s = 1/n_T$. Segment end points are denoted $\mathbf{X}^{[n]}(t) := \mathbf{X}(s^{[n]}, t)$, at arclength $s^{[n]} := \Delta s$, for $n = 0, \ldots, n_T$ (where $\mathbf{X}^{[0]}$ is the point joining the head and flagellum). The angle joining $\mathbf{X}^{[n-1]}$ to $\mathbf{X}^{[n]}$ is denoted $\tilde{\theta}^{[n]}$ for $n = 1, \ldots, n_T$. Approximating the force density exerted by the flagellum on the fluid as piecewise constant, we write $\mathbf{f}(s,t) \approx \tilde{\mathbf{f}}^{[n]} = \mathbf{f}(\tilde{s}^{[n]}, t)$, with $\tilde{s}^{[n]} = (n - 1/2)\Delta s$ for $n = 1, \ldots, n_T$. The active moment is approximated as piecewise linear along each segment, with $m^{[n]} := m(s^{[n]}, t)$, $n = 0, \ldots, n_M - 1$ denoting the active moment at the actively bending segment joints. Here, $n_M = 57 < n_T$ has been chosen to represent the approximate 95% of the flagellum that actively bends.

#### 1.3 Discretised system

The spatially discretised equations can then be written, for the force- and moment-free conditions as

$$\sum_{n=1}^{n_H} \varphi_j^{[n]} \sum_{m=1}^{n_Q} \delta_{ij} \nu^{[m,n]} + \sum_{n=1}^{n_T} \Delta s\, \tilde{f}_i^{[n]} = 0, \quad i = 1, 2; \tag{S1}$$

$$\delta_{i3} \sum_{n=1}^{n_H} \epsilon_{ijk} \left[ Y_j^{[n]} - X_j^{[1]} \right] \varphi_k^{[n]} \sum_{m=1}^{n_Q} \delta_{ij} \nu\,[m,n] + \sum_{n=1}^{n_T} \Delta s \epsilon_{ijk} \left[ \tilde{X}_j^{[n]} - X_j^{[1]} \right] \tilde{f}_k^{[n]} = 0; \tag{S2}$$

for the elastic behaviour of the flagellum as

$$E\left(s^{[n]}\right)\left[\frac{\tilde{\theta}^{[n+1]}-\tilde{\theta}^{[n]}}{\Delta s}\right]-\mathcal{M}\Delta s\left[\frac{1}{2}m^{[n]}+\sum_{i=n+1}^{n_M-1}m^{[i]}\right]=-\mathcal{S}^4\mathbf{e}_3\cdot\sum_{i=n}^{n_T-1}\Delta s\left[\tilde{\mathbf{X}}^{[i+1]}-\mathbf{X}^{[n]}\right]\times\tilde{\mathbf{f}}^{[i+1]},\ n=1,\ldots,n_M; \tag{S3}$$

$$E\left(s^{[n]}\right)\left[\frac{\tilde{\theta}^{[n+1]}-\tilde{\theta}^{[n]}}{\Delta s}\right]=-\mathcal{S}^4\mathbf{e}_3\cdot\sum_{i=n}^{n_T-1}\Delta s\left[\tilde{\mathbf{X}}^{[i+1]}-\mathbf{X}^{[n]}\right]\times\tilde{\mathbf{f}}^{[i+1]},\ n=n_M,\ldots,n_T-1; \tag{S4}$$

$$E\left(s^{[0]}\right)\left[\frac{\tilde{\theta}^{[1]}-\phi}{\Delta s}\right]-\mathcal{M}\Delta s\left[\frac{1}{2}m^{[0]}+\sum_{i=1}^{n_M-1}m^{[i]}\right]=-\mathcal{S}^4\mathbf{e}_3\cdot\sum_{i=0}^{n_T-1}\Delta s\left[\tilde{\mathbf{X}}^{[i+1]}-\mathbf{X}^{[0]}\right]\times\tilde{\mathbf{f}}^{[i+1]}; \tag{S5}$$

for the hydrodynamic equations evaluated for points on the flagellum, with $n=1,\ldots,n_T$, as

$$\begin{aligned}\dot{X}_i^{[0]}+\mathbf{e}_i\cdot&\left[\frac{\Delta s}{2}\dot{\tilde{\theta}}^{[n]}\left[-\sin\tilde{\theta}^{[n]},\cos\tilde{\theta}^{[n]},0\right]^T+\sum_{k=1}^{n-1}\Delta s\,\dot{\tilde{\theta}}^{[k]}\left[-\sin\tilde{\theta}^{[k]},\cos\tilde{\theta}^{[k]},0\right]^T\right]\\&=\sum_{k=1}^{n_H}\varphi_j^{[k]}\sum_{l=1}^{n_Q}S_{ij}^{\varepsilon}\left(\tilde{\mathbf{X}}^{[n]},\hat{\mathbf{Y}}^{[l]}\right)\nu^{[k,l]}+\sum_{k=1}^{n_T}I_{ij}^{[k]}\left(\tilde{\mathbf{X}}^{[n]},t;\Delta s,\varepsilon\right)\tilde{f}_j^{[k]},\ i=1,2,3;\end{aligned} \tag{S6}$$

where the integral

$$I_{ij}^{[k]}\left(\chi,t;\Delta s,\varepsilon\right)=\int_{(k-1)\Delta s}^{k\Delta s}S_{ij}^{\varepsilon}\left(\chi,\tilde{\mathbf{X}}^{[k]}+\left(s-\tilde{s}^{[k]}\right)\left[\cos\tilde{\theta}^{[k]},\sin\tilde{\theta}^{[k]},0\right]^T\right)\mathrm{d}s, \tag{S7}$$

is calculated analytically following Refs. [7, 8, 1]; the hydrodynamic equations evaluated for points on the head, with $n=1,\ldots,n_H$, are written as

$$\mathbf{e}_i\cdot\left[\dot{\mathbf{X}}^{[0]}+\dot{\phi}\,\mathbf{e}_3\times\left[\mathbf{Y}^{[n]}-\mathbf{X}^{[0]}\right]\right]=\sum_{k=1}^{n_H}\varphi_j^{[k]}\sum_{l=1}^{n_Q}S_{ij}^{\varepsilon}\left(\mathbf{Y}^{[n]},\hat{\mathbf{Y}}^{[l]}\right)\nu^{[k,l]}+\sum_{k=1}^{n_T}I_{ij}^{[k]}\left(\mathbf{Y}^{[n]},t;\Delta s,\varepsilon\right)\tilde{f}_j^{[k]},\ i=1,2,3; \tag{S8}$$

for the rate equation governing the active moment, for $n=1,\ldots,n_M$, as

$$\dot{m}^{[n]}=-\left(m^{[n]}-2\,\mathrm{sgn}\left(m^{[n]}\right)\right)\mathcal{H}\left(\kappa_c-\,\mathrm{sgn}\left(m^{[n]}\right)\frac{\tilde{\theta}^{[n+1]}-\tilde{\theta}^{[n]}}{\Delta s}\right)-\mathrm{sgn}\left(m^{[n]}\right). \tag{S9}$$

The spatially discrete system can then be written in the form of a non-linear autonomous initial value problem,

$$\dot{\mathbf{z}}=\mathcal{F}(\mathbf{z}),\qquad \boldsymbol{z}(0)=\boldsymbol{z}_0, \tag{S10}$$

where $\dot{\mathbf{z}}=\partial\boldsymbol{z}/\partial t$, with $\mathbf{z}(t):=\left[\mathbf{X}^{[1]}(t),\phi(t),\tilde{\theta}^{[1]},\ldots,\tilde{\theta}^{[n_T]},\tilde{m}^{[1]}(t),\ldots,\tilde{m}^{[n_M]}(t)\right]^T$. The system (S10) can be augmented at each time point to include the unknown force densities as

$$A\begin{bmatrix}\dot{\mathbf{z}}\\ \mathbf{f}\\ \boldsymbol{\varphi}\end{bmatrix}=\mathbf{b}, \tag{S11}$$

with

$$A=\left[\begin{array}{c|c|c}0 & A_E & 0\\ \hline A_K & A_H & 0\\ \hline 0 & 0 & A_M\end{array}\right]. \tag{S12}$$

The matrix blocks of $A$ $(A_E,A_K,A_H,A_M)$ encode the elastodynamic, force- and moment-free equations (Eqs. S1-S5); the kinematic and hydrodynamic equations (Eqs. S6-S8); and active moment equations ((S9)) respectively.

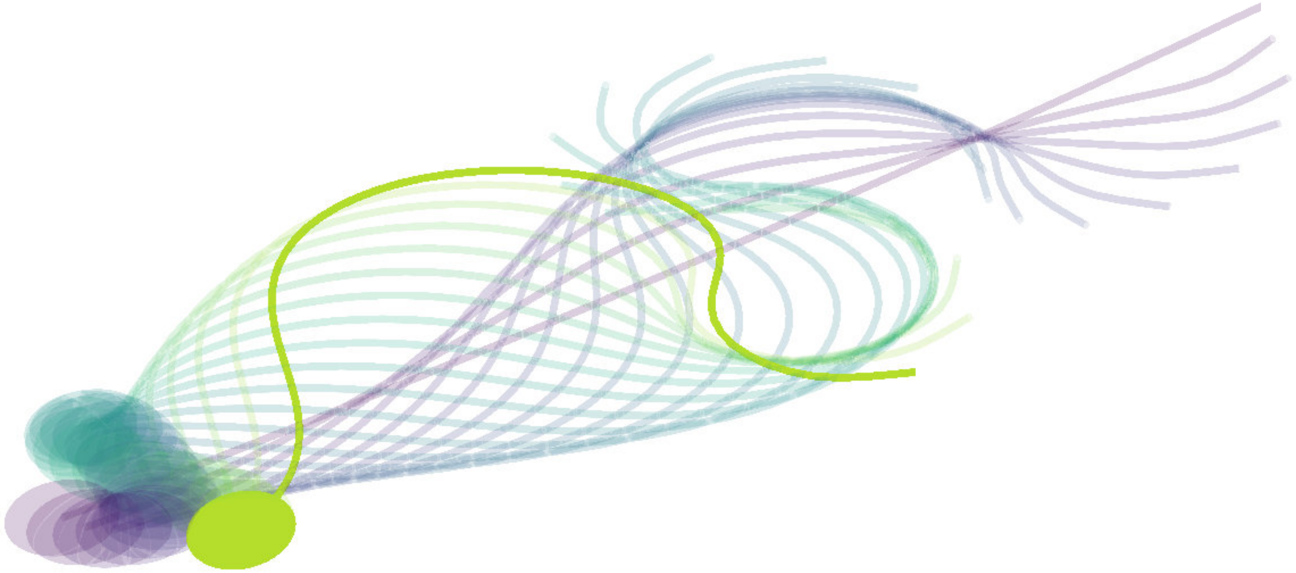

Figure 1: Evolution of the flagellar beat from the initial low-amplitude parabola.

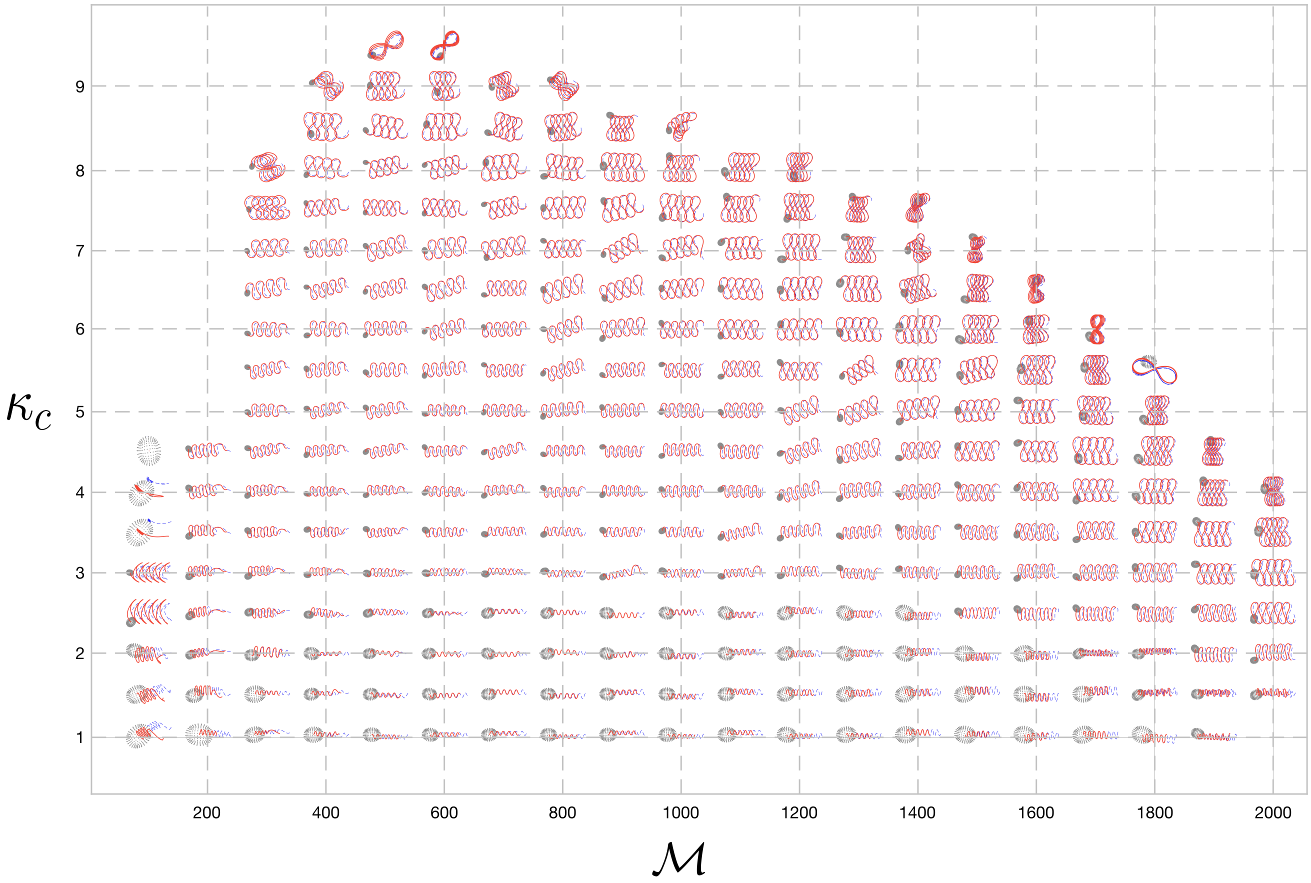


Figure 2: Sperm tracks from simulations plotted across the regulatory parameter subspace $(\mathcal{M}, \kappa_c)$. The active moment $(\mathcal{M})$ and critical curvature $(\kappa_c)$ parameters are varied with $(\mathcal{S}, \rho) = (18, 36.4)$. Parameter choices with missing entries (e.g. $(\mathcal{M}, \kappa_c) = (100, 5)$) are unable to generate a beat. Red solid lines show the path traced about by the centroid of the head, while the blue dashed curves show the path of the head/flagellum join.

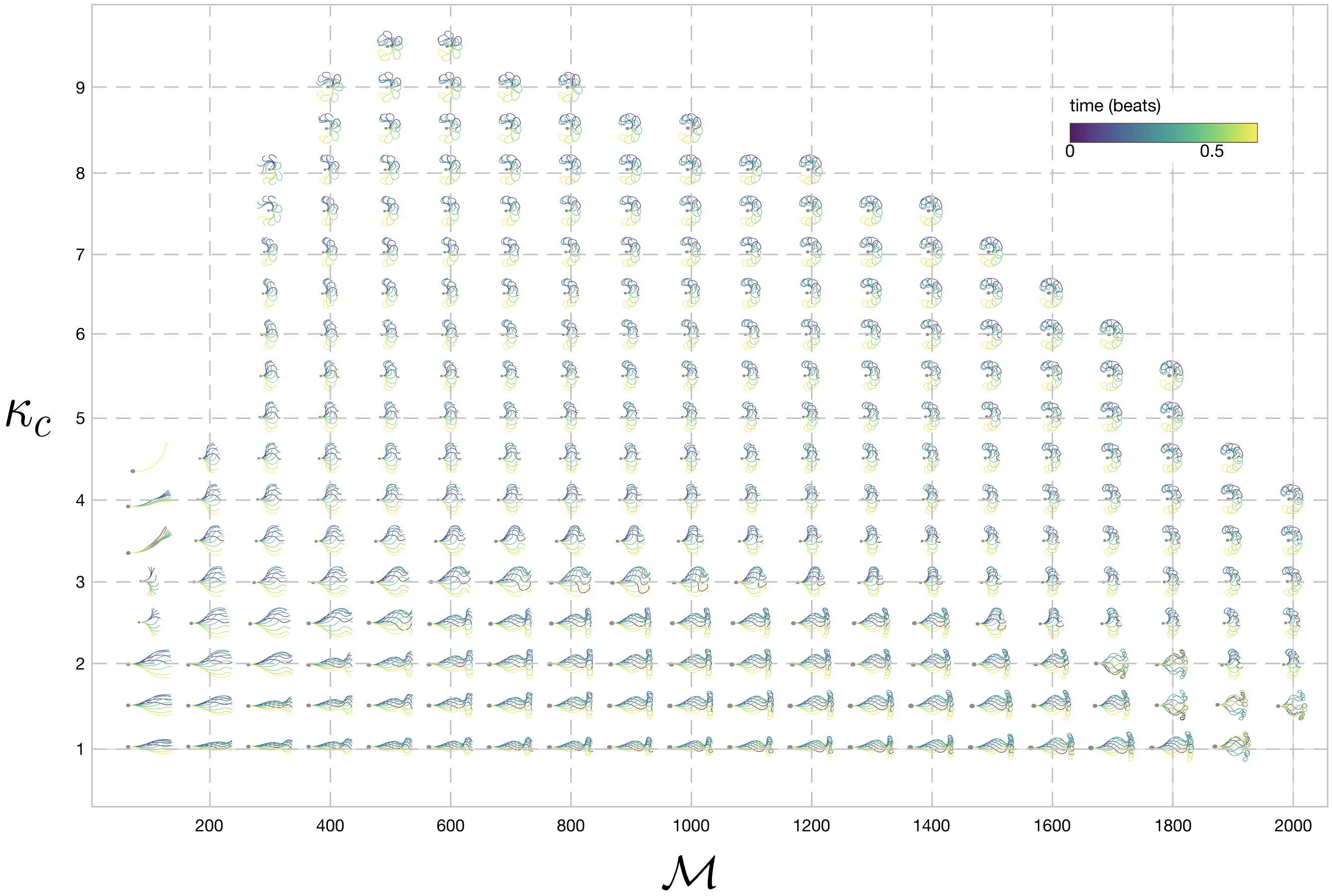


Figure 3: Sperm waveforms from simulations plotted across the regulatory parameter subspace $(\mathcal{M}, \kappa_c)$. The active moment $(\mathcal{M})$ and critical curvature $(\kappa_c)$ parameters are varied with $(\mathcal{S}, \rho) = (18, 36.4)$. Parameter choices with missing entries (e.g. $(\mathcal{M}, \kappa_c) = (100, 5)$) are unable to generate a beat.

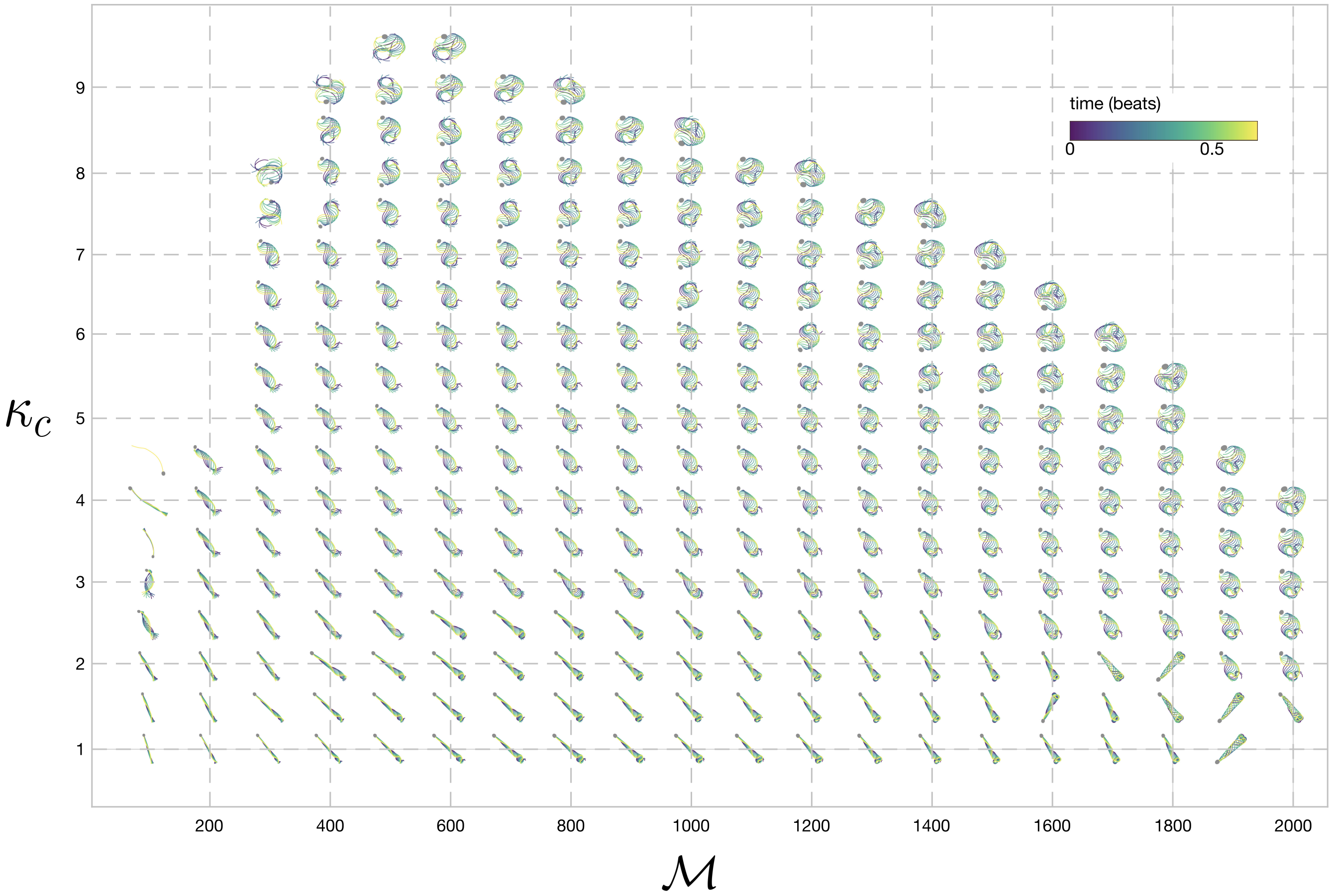


Figure 4: Sperm flagellar time-lapse from simulations plotted across the regulatory parameter subspace $(\mathcal{M}, \kappa_c)$. The active moment $(\mathcal{M})$ and critical curvature $(\kappa_c)$ parameters are varied with $(\mathcal{S}, \rho) = (18, 36.4)$. Parameter choices with missing entries (e.g. $(\mathcal{M}, \kappa_c) = (100, 5)$) are unable to generate a beat.

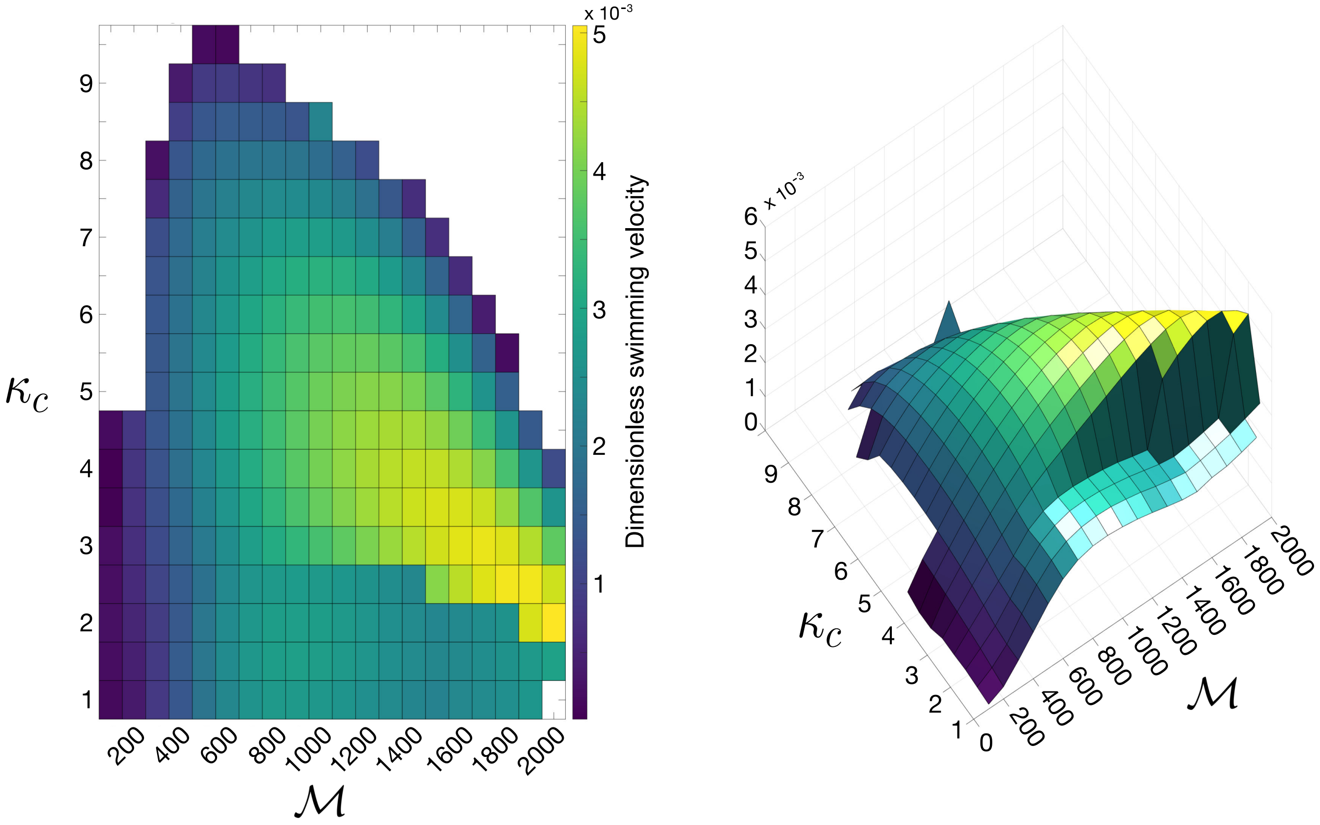


Figure 5: Dimensionless swimming velocity of sperm simulations plotted across the regulatory parameter subspace $(\mathcal{M}, \kappa_c)$. The active moment $(\mathcal{M})$ and critical curvature $(\kappa_c)$ parameters are varied with $(\mathcal{S}, \rho) = (18, 36.4)$. Parameter choices with missing entries (e.g. $(\mathcal{M}, \kappa_c) = (100, 5)$) are unable to generate a beat.

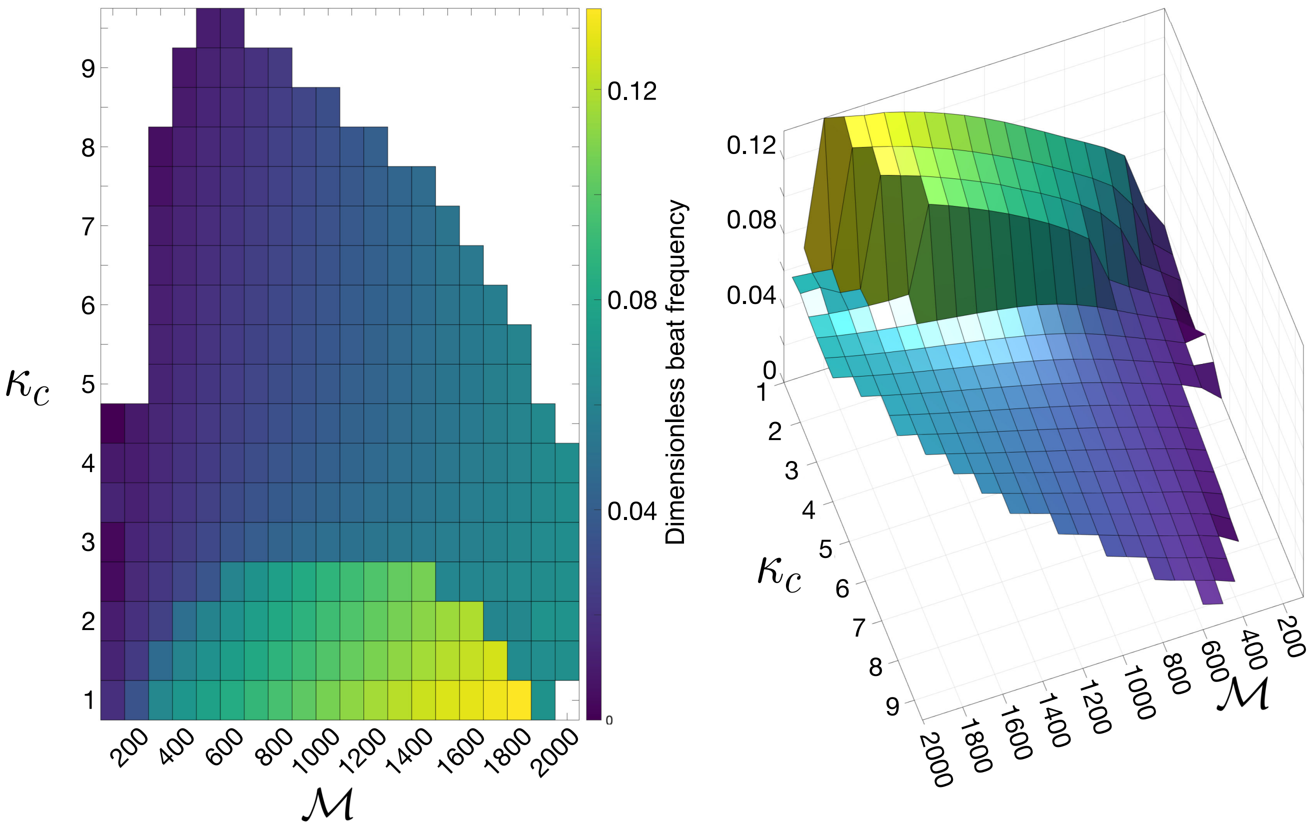


Figure 6: Dimensionless flagellar beat frequency of sperm simulations plotted across the regulatory parameter subspace $(\mathcal{M}, \kappa_c)$. The active moment $(\mathcal{M})$ and critical curvature $(\kappa_c)$ parameters are varied with $(\mathcal{S}, \rho) = (18, 36.4)$. Parameter choices with missing entries (e.g. $(\mathcal{M}, \kappa_c) = (100, 5)$) are unable to generate a beat.
